# dipwmsearch: a python package for searching di-PWM motifs

**DOI:** 10.1101/2022.11.08.515647

**Authors:** Marie Mille, Julie Ripoll, Bastien Cazaux, Eric Rivals

## Abstract

**Motivation:** Seeking probabilistic motifs in a sequence is a common task to annotate putative transcription factor binding sites (TFBS). Useful motif representations include Position Weight Matrices (PWMs), dinucleotidic PWMs (di-PWMs), and Hidden Markov Models (HMMs). Dinucleotidic PWMs combine the simplicity of PWMs – a matrix form and a cumulative scoring function –, but also incoporate dependency between adjacent positions in the motif (unlike PWMs which disregard any dependency). For instance, to represent binding sites, the HOCOMOCO database provides di-PWM motifs derived from experimental data. Currently, two programs, SPRy-SARUS and MOODS, can search for di-PWMs in sequences.

**Results:** We propose a Python package, *dipwmsearch*, which provides an original and efficient algorithm for this task (it first enumerates matching words for the di-PWM, and then search them at once in the sequence even if it contains IUPAC codes). The user benefits from an easy installation via *Pypi* or *conda*, a documented Python interface, and reusable example scripts that smooth the use of di-PWMs.

**Availability and Implementation:** *dipwmsearch* is available at https://pypi.org/project/dipwmsearch/ and https://gite.lirmm.fr/rivals/dipwmsearch/ under Cecill license.

## 1 Introduction

Protein binding sites on nucleic acids (DNA or RNA) share similar, but not identical sequences. The collection of sequences of such binding sites, which in practice is a set of sequences (of identical length), are summarized and represented as a probabilistic motif. Often only a few positions within such sequences are conserved across a majority of binding sites. Even at a conserved position, when the collection is large enough, alternative nucleotides occur. Hence, for each position of the binding site, it is convenient to summarize its variability as the probability of each nucleotide to occur at this position. The probabilities are estimated from the frequencies of nucleotides at that position in the collection. This explains why the first and most popular probabilistic motif representation is the Position Weight Matrix (PWM) [8]. A PWM is a matrix containing the weight or score of each nucleotide at each position of the sequence alignment: the weights that are log-odd scores of the nucleotide probabilities at each position. Numerous search algorithms are available for PWMs [3, 2, 7]. However, in a PWM positions are entirely independent one of another; but in reality neighboring positions are constrained for they influence the shape of DNA, or the propensity to undergo epigenetic modifications, and hence the binding of the protein. Hence, a more complex representation for probabilistic motifs that accounts for local position dependencies was proposed: dinucleotidic PWM (di-PWM) [5]. At each position, one records the frequency of all 16 possible dinucleotides (instead of 4 nucleotides for PWM). A di-PWM and score computation of a word is illustrated Figure 1a.

**Figure 1:**
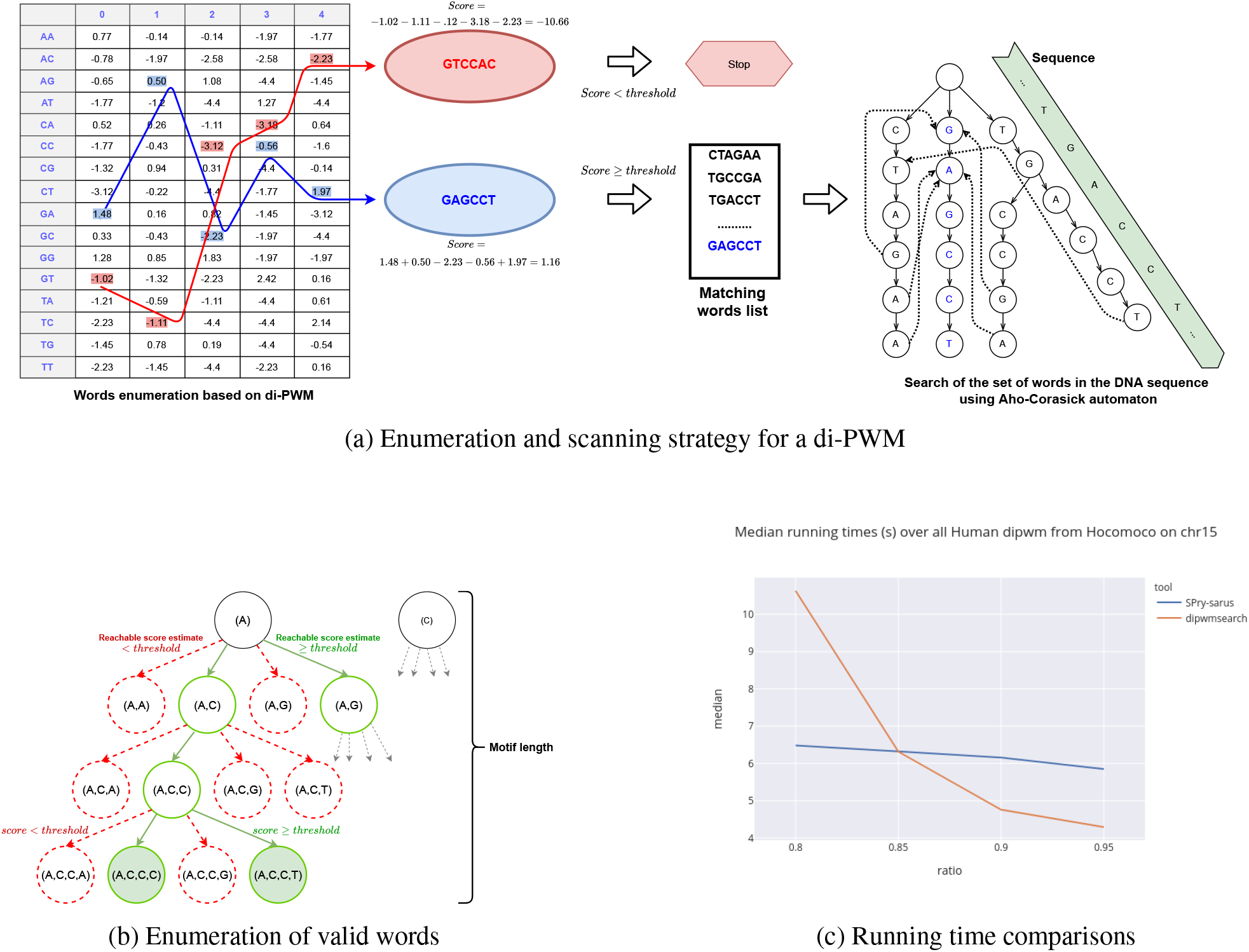
**(a)** Enumeration and scanning strategy for a di-PWM. Left part shows how the score of two words are computated by summing the score of their five dinucleotides. If the score lies above the threshold, the word is a valid word and is added to the list for later search. Right part: we build an Aho-Corasick automaton with all valid words in the list, then use the automaton to scan the sequence. **(b)** Illustration of the branch and bound strategy for the enumeration procedure. We build a trie for words starting with letter *A*, and explore it in Depth-First manner. As soon as a prefix cannot give rise to a valid word, which is determined using the LookAheadMatrix (LAM), we cut the corresponding branch. Only valid words generate a leaf in the trie. **(c)** Comparison of SPRy-SARUS and dipwmsearch for searching all Human di-PWMs from HOCOMOCO on Human chromosome 15. The plot shows the median running time over all di-PWMs.

For instance, the *HOCOMOCO* database (v11) stores di-PWMs of binding motifs of Human and Mouse transcription factors (292 and 257, respectively), which were directly computed from experimental ChIP-Seq data [6]. For detecting new binding sites, it was shown that di-PWMs provide enhance sensitivity compared to classical PWMs [5]. Results of DREAM-ENCODE challenge from 2017 tend to confirm this observation. To our knowledge, only *SPRy-SARUS* and *MOODS* can find occurrences of di-PWMs in long sequences. *SPRy-SARUS* is an efficient stand alone Java program to search for di-PWMs, which is coupled with *MoLoTool*, a webtool allowing visual inspection of occurrences in short sequences [6]. *MOODS*, a tool to search for PSSM/PWM, can also handle di-PWMs [4] (v3 available as C++ code and python package). Both adopt a window scanning strategy, while dipwmsearch uses an enumeration strategy. Hence, we provide a Python package for searching di-PWM in sequences: it can be easily installed via *conda* and offers several functions that can be used in Python programs. We designed a novel search algorithm that differs from previous approaches. Running time comparisons demonstrate that our algorithm is on par with SPRy-SARUS in practice.

## 2 Search algorithm

Our package provides distinct search algorithms: an optimized scanning algorithm (OS), an enumeration based algorithm for full di-PWM (FE), and core enumeration based algorithm (CE). For place sake, we described only CE algorithm for it is the most efficient. Algorithm viewpoint, we aim at proposing new approaches capable of searching for matches in long sequences, with limited amount of memory. Before explaining the algorithm, we give some rationale for our approach. Let us consider that in a text *T*, we seek a di-PWM *P* of size σ^2 ×^(*m*−1) (for a motif of length *m*) and with a score threshold *t*. An entry *P*[αβ, *i*] gives the score of dinucleotide αβ at position *i* in the motif. Remark that for any interval [*i*.. *j*] with 1 *≤ i < j < m*, the restriction of *P* to this interval of positions, which we denote by *P*[][*i*.. *j*], is a smaller di-PWM of length *j* −*i* + 1.

### Traditional scanning algorithm and the enumeration strategy

In a traditional scanning algorithm, one considers each possible window of length *m* in *T* and computes its score according to *P*. It takes *O*(*m* × |*T*|) time, which is quadratic. The scanning approach implies redundant computation (for instance when processing identical or similar substrings whose score are too low) and often is inefficient. A classical speedup trick uses the LookAheadTable to stop the score computation after viewing only a prefix of the current window [2]; it does not improve the worst case complexity.

An alternative is to first enumerate all words of length *m* that match *P* with a score *> t*, which we call *valid words*, and then to search the valid words in *T* using an Aho-Corasick (AC) automaton [1] (or any other algorithm that solves the Set Pattern Matching problem). We implemented this in the enumeration based algorithm for full di-PWM (FE). We call this global idea the *enumeration strategy*; it concentrates the complexity in the enumeration phase, and makes the scanning efficient because it seeks `only exact` matches of the valid words – the scanning phase does not compute any score. Nicely, building the Aho-Corasick automaton takes linear time in cumulated length of valid words, and scanning *T* with it takes linear time in |*T* |.

### Efficiency conditions for the enumeration strategy

To be efficient, the enumeration strategy needs 1/ a fast enumeration algorithm, 2/ a set of valid words that is small enough for the AC automaton to fit in memory (i.e., remain fast to build). Below, we exhibit a enumeration algorithm takes linear time in the output size, which resolves the first condition. However, the number of valid words depends on the selectivity of the di-PWM *P* with threshold *t*. The least selective position is when the scores of all 16 dinucleotides are equal. It turns out that some di-PWMs from HOCOMOCO contain positions that are not selective, i.e., in which the scores are almost equally distributed. A closer examination shows in such di-PWMs non selective positions often occur in intervals of successive positions (see Figure 1 in Supplementary Material). For such an interval of say *f* positions, ify we consider *P*′ the restriction of *P* to this interval, almost any word of length *f* + 1 is a valid word for the di-PWM *P*′. This may lead to an explosion of valid words for the full di-PWM *P*.

We propose to identify selective and non-selective positions by considering the standard deviation of their scores: a large deviation means a selective position. To avoid cases with huge set of valid words, we propose to restrict *P* to an interval of selective positions, which we term the *core*. We proceed as follows: first we compute the standard deviation of scores for all positions, then we select, by exhaustive search among all possible intervals of length at least 10, the interval with the largest average standard deviation. This interval determines the core (which is a smaller di-PWM).

### Enumeration of valid words: B&B approach and LAM

For a full di-PWM, we propose an algorithm that explores a trie data structure of valid words using a Branch-and-Bound approach (see Figure 1b). We build a trie that spells out prefixes of potential valid words, one letter at a time. After each letter, assume the current prefix has length *k*, we compute the partial score for this prefix. Then, we check the score for the best possible suffix of length *m* − *k* in an additional matrix called the LAM. If the sum of prefix and suffix scores does not reach the threshold *t*, then extensions of the current branch of the trie are unnecessary. The LookAheadMatrix (or LAM for short) is a precomputed σ× (*m*− 1) matrix that depends only on *P*. For a position *i* in *P* and a symbol α, the *LAM*[α, *i*] stores the best score for a suffix starting with symbol α at position *i*. Algorithm **??** computes the LAM in *O*(σ^2^ × (*m*− 1)) time. The LAM has a crucial property: for any stored score value in the LAM, there exists a word that realises this score. This ensures that only branches of the trie corresponding to valid words are fully built by the enumeration algorithm. Moreover, the amount of computation spent between two successive valid words is bounded by 2*m*, which implies that our algorithm takes linear time in the output size.

Note that a pendant matrix to the LAM can be built symmetrically to compute the best scores of prefixes of *P*. We call this matrix, the LookBackMatrix or LBM.

After enumeration, in the search step, the set of valid words for *P* are searched for in *T* using an Aho-Corasick automaton [1].

### Adapting the enumeration strategy and search with the core

The enumeration algorithm and search phase must be adapted to use the core instead of *P*. Assume the core, denoted by *Q*, starts at position *k* + 1 in the motif and has length *h* − 1. We must enumerate words of length *h* for *Q* that are substrings of valid words of length *m* for *P*. We run the branch & bound algorithm described above to spell out words of length *h* according to *Q*, but we cannot select them on their own score (which is a sum only over *h* − 1 positions!). We must use the score of a prefix of *P*, not a prefix of *Q*. Assume the current prefix *w* starts with letter α at position *k* + 1 and ends with letter β; as score, we use score(*xwy*), where *x* is a highest scoring valid prefix for *P* of length *k* ending with letter α, and *y* a highest scoring suffix of length *m*− *k*−|*w*| + 1 starting with letter β, and such that |*xwy*| = *m*. The idea behind is that *xwy* is the best possible word of length *m* with substring *w* (at positions [*k* + 1, *k* + |*w*| + 1]). The constraints on the letters are implied by the fact that successive positions of a di-PWM score overlapping dinucleotides. We use the LAM to get the contribution of *y* to this score (without knowing *y*) and we use the LBM to get that of *x* (without knowing *x*). The algorithm outputs the set of all words *w* of length *h* that occur in at least one valid word for *P* (at position [*k* + 1, *k* + *h* + 1]).

The search phase builds an AC automaton with this set and scan *T* with it. Each time a match is found, say at position *i*, it computes the score of the window of *T* between positions (*i*−*k* + 1) and (*i*−*k* + *m*). If the scores reaches the threshold *t* it reports a match of *P* at position (*i*−*k* + 1) in *T*, and its score.

## 3 Results

We compare the core motif algorithm to SPRy-SARUS in terms of efficiency by searching each Human di-PWM from HOCOMOCO on Human chromosomes 3 and 15 for four different score ratios (0.8, 0.85, 0.90, and 0.95). First, this confirms that core algorithm is able to search any di-PWM with reasonable score ratios. Figure 1c displays the median search time over all di-PWMs for both tools and shows first, that dipwmsearch offers affordable runtimes whatever the ratio, and second that dipwmsearch takes longer times than SPRy-SARUS for ratio 0.80, while it is as efficient or faster for larger ratios.

## 4 Conclusion

Our Python package, *dipwmsearch*, provides an easy use of an efficient procedure to search di-PWM in nucleotidic sequences, through a set of documented snippets. It offers practical advantages compared to an existing solution (like processing IUPAC codes, or an adaptable output – see Supplementary Material) and can be enhanced by combining it with other Python packages (e.g., for processing compressed sequence files). Most of all, installation is straighforward using *pypi* or *conda*. In addition, we presented an original, enumeration based search algorithm that can handle di-PWMs even if they contain non selective positions – which seems novel. Coping with non selective positions was necessary to make search effective for some di-PWMs, which questions their information content, and in turn their construction process. Examining the set of valid words and their occurrences could help determining if a di-PWM model is truely well suited.

Several perspectives come to mind. First, once enumerated, the set of valid words can stored in a file and reused for other searches. Second, the search phase can be streamlined by using a precomputed index of the search sequence to find valid words, which would enable a web application to support numerous di-PWM searches.

## Supporting information

Supplementary Material (5 pages, 6 sections, and references)

