## Supplementary Material (5 pages, 6 sections, and references) for "dipwmsearch: a python package for searching di-PWM motifs"

### dipwmsearch: a python package for searching di-PWM motifs – Supplementary Material

November 8, 2022

**Abstract**

**Motivation** Seeking probabilistic motifs in a sequence is a common task to annotate putative transcription factor binding sites (TFBS). Useful motif representations include Position Weight Matrices (PWMs), dinucleotidic PWMs (di-PWMs), and Hidden Markov Models (HMMs). Dinucleotidic PWMs combine the simplicity of PWMs – a matrix form and a cumulative scoring function –, but also incoporate dependency between adjacent positions in the motif (unlike PWMs which disregard any dependency). For instance, to represent binding sites, the HOCOMOCO database provides di-PWM motifs derived from experimental data. Currently, two programs, SPRy-SARUS and MOODS, can search for di-PWMs in sequences.

**Results** We propose a Python package, *dipwmsearch*, which provides an original and efficient algorithm for this task (it first enumerates matching words for the di-PWM, and then search them at once in the sequence even if it contains IUPAC codes). The user benefits from an easy installation via *Pypi* or *conda*, a documented Python interface, and reusable example scripts that smooth the use of di-PWMs.

**Availability and Implementation**: *dipwmsearch* is available at https://pypi.org/project/dipwmsearch/ and https://gite.lirmm.fr/rivals/dipwmsearch/ under Cecill license.

#### 1 Access to the packages, the source code and the documentation.

1. Python package: `https://pypi.org/project/dipwmsearch/`
2. Documentation: `https://rivals.lirmm.net/dipwmsearch/`
3. Conda package: `https://anaconda.org/atgc-montpellier/dipwmsearch`
4. Source code: `https://gite.lirmm.fr/rivals/dipwmsearch`
5. Contact:

#### 2 Practical comparison of SPRy-SARUS, MOODS, and dipwmsearch

We summarize some features between the tools SPRy-SARUS (url), MOODS [1], and dipwmsearch [3], in the table below.

| **Feature** | SPRy-SARUS | MOODS v3 | dipwmsearch |
|---|---|---|---|
| **conda installation** | n | y | y |
| **pypi installation** | n | y | y |
| **IUPAC code (except N)** | blocking | y | y |
| **API programming library** | n | y | y |
| **output matching word** | n | y | y |
| **compressed sequence file** | n | y | y (via gzip/bzip python packages) |
| **multi FASTA** | y | y | y (example) |

#### 3 Simplicity of use: an example of python code

To illustrate how easily one can use dipwmsearch in python, we provide an example of code for searching one di-PWM motif in a FASTA sequence for a given threshold, exactly as SPRy-SARUS would perform it. This script takes three arguments on the command line: first, the file containing the di-PWM, second, the file containing the FASTA formatted sequence, third the threshold ratio as a real value.

```
import sys, os
import dipwmsearch as ds
from Bio import SeqIO
from Bio.Seq import Seq

pathSeq = sys.argv[1]
pathDiPwm = sys.argv[2]
threshold = float(sys.argv[3])


# read diPWM, sequence
diP = ds.create_diPwm(pathDiPwm)
file = open(pathSeq)
seqRecord = SeqIO.read(file, "fasta")

mySeq = str(seqRecord.seq.upper())
seq = mySeq.translate(mySeq.maketrans("NMRWSYKVHDB","GGGGGGGGGGG"))

# print(threshold)

# Block optimized search
number_matches_block_opt = 0
# 1st loop to search valid words on Watson strand
for i, word, score in ds.search_block_optimized(diP, seq, threshold):
    number_matches_block_opt += 1

# reverse complement sequence
seq_rev = seq.translate(seq.maketrans("ACGT","TGCA"))
seq_rev = seq_rev[::-1]

# 2nd loop to search valid words on Crick strand
for i, word, score in ds.search_block_optimized(diP, seq_rev, threshold):
    number_matches_block_opt += 1

dipwm_name = os.path.split(pathDiPwm)[1].split(".")[0]
print(f'{dipwm_name}\t{threshold}\t{number_matches_block_opt:,}')
```

#### 4 LAM: definition and algorithm

**Definition 4.1 (*LookAheadMatrix* for di-PWM)** *The* LookAheadMatrix *$M$ of a di-PWM $P$ of a motif of length $m$ on an alphabet $\Sigma$ of size $\sigma$ is a matrix of size taille $\sigma \times (m-1)$, where for any $i$ such that $0 \leq i \leq m-2$ and for any $d \in \Sigma$:*

$$M[d,i] := \begin{cases} \max_{b \in \Sigma} P[db,i], & \text{if } i = m-2 \\ \max_{b \in \Sigma} (P[db,i] + M[b,i+1]), & \text{if } 0 \leq i < m-2 \end{cases}$$

**Algorithm 1:** MakeLookAheadMatrix: compute the LookAheadMatrix of a di-PWM $P$.

**Input:** Alphabet $\Sigma$ of size $\sigma$,di-PWM matrix $P$ of size $\sigma^2 \times (m-1)$, for a motif of length $m$
**Output:** LookAheadMatrix $M$ of size $\sigma \times (m-1)$

$M \leftarrow$ initialized with $-\infty$
**for** $d \in \{0,\dots,\sigma-1\}$ **do**
  $max \leftarrow -\infty$
  **for** $b \in \{0,\dots,\sigma-1\}$ **do**
    $score \leftarrow P[db, m-2]$
    **if** $score > max$ **then** $max \leftarrow score$
  $M[d, m-2] \leftarrow max$
**for** $i \in \{m-3,\dots,0\}$ **do**
  **for** $d \in \{0,\dots,\sigma-1\}$ **do**
    $max \leftarrow -\infty$
    **for** $b \in \{0,\dots,\sigma-1\}$ **do**
      $score \leftarrow P[db, i] + M[b, i+1]$
      **if** $score > max$ **then** $max \leftarrow score$
    $M[d, i] \leftarrow max$
**return** $M$

### 5 Non selective positions and core

An example of di-PWM with non selective positions, for GATA2 transcription factor, is shown Figure 1. The height of the letters at each motif position in the LOGO above the line, indicate the information content of the corresponding nucleotide at this position. The letters first 12 positions are nearly invisible compared to those between position 13 until 22.

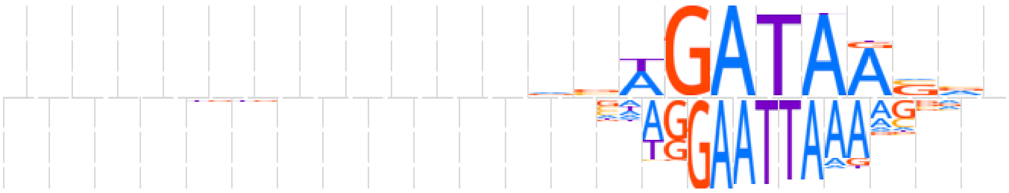

Figure 1: LOGO representation of the Human di-PWM for the binding site motif of transcription factor GATA2. The motif 23 bp long and its consensus sequence is *nnnnnnnnnnnnvvWGATAASvn*. The interval with the first 12 positions at 5' end contains non selective positions (with little information content), as is the last 3' end position. Hence, the core is restriction to interval [13, 22] of positions.

### 6 Protocol for comparing running times

Currently, it is not possible to search for di-PWM using MOODS v3: MOODS cannot read the format for di-PWM matrices containing scores (an issue was sent on the github repository on Sep 22, 2022). This is why MOODS was not used in the comparisons.
Comparisons between dipwmsearch and SPRy-SARUS were performed as follows, using two bash scripts on a Linux system. Technical information are summarised below.
For both chromosome sequences (15 and 3), for all Human di-PWM motifs of HOCOMOCO database [2], for four ratios (0.8, 0.85, 0.9, 0.95), we recorded the running time of dipwmsearch and SPRy-SARUS using

the `time` command. Each search for a di-PWM was performed in a distinct execution of the tools, implying that each time the target sequence and the di-PWM were loaded, the enumeration of valid words and then the scanning of the sequence performed, before storing the results in an output file on the disk. We did not take advantage of the possibility for dip to search for a multi FASTA file containing both sequences at once (which would have avoided to redo the enumeration step).
For dipwmsearch, we used a python script that calls the procedure `search_block_optimized` for searching the di-PWM first on the Watson strand and then on the Crick strand. The time was measured using the `time` command from Python package `time`. For SPRy-SARUS, we recorded the time taken by program in user mode, using the `/usr/bin/time` command with option `-f "%U"`. Both tools were run using a single thread; despite this, it occurs that the JAVA machine running SPRy-SARUS used more than 100% of the CPU.

##### Technical information

The last version of SPRy-SARUS (release: 2.0.2) was obtained from its github repository https://github.com/autosome-ru/sarus.
**Linux version**
Linux 5.11.0-40-generic #44~20.04.2-Ubuntu SMP Tue Oct 26 18:07:44 UTC 2021 $x86_{64}$ $x86_{64}$ $x86_{64}$ GNU/Linux
**Java version**

- openjdk version "10.0.2" 2018-07-17
- OpenJDK Runtime Environment Zulu10.3+5 (build 10.0.2+13)
- OpenJDK 64-Bit Server VM Zulu10.3+5 (build 10.0.2+13, mixed mode)

##### Variation of running time with respect to the HOCOMOCO di-PWM

The summary of results reported in the article gives the median running times for all HOCOMOCO di-PWM motifs. As mentioned above, the di-PWMs of HOCOMOCO vary in information content and thus in selectivity. This implies that the number of valid words strongly depends on the motif and the ratio, and hence, so do the enumeration and overall running times.
The Figures 2 provides a view on the variability of the running times in function of the di-PWM and ratio, for both SPRy-SARUS (left plot) and dipwmsearch (right plot). We report the running times for searching on Human chromosomes 15 and 3. Note that because SPRy-SARUS can only process sequences containing A, C, G, T (but no other IUPAC codes), the sequence of chromosome 3 was slightly modified to replace undetermined positions by one standard nucleotide (before feeding SPRy-SARUS). For dipwmsearch, we used the normal chromosome 3 sequence with IUPAC codes.
The variation of running times in comparison to the median running time is higher for dipwmsearch than for SPRy-SARUS. For dipwmsearch, it depends on the matrix and ratio, while SPRy-SARUS times depend mostly on the sequence length. However, the overal running times of dipwmsearch remain fast enough for practical uses.

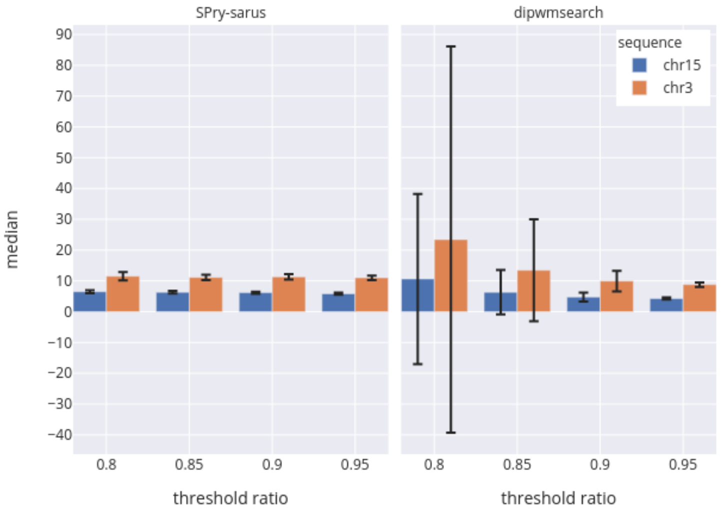


Figure 2: Median running times and standard deviations of both dipwmsearch and SPRy-SARUS for searching each Human di-PWMs from HOCOMOCO on Human chromosomes 15 and 3.
